# Evolution of exploitative interactions during diversification in *Bacillus subtilis* biofilms

**DOI:** 10.1101/173476

**Authors:** Anna Dragoš, Nivedha Lakshmanan, Marivic Martin, Balázs Horváth, Gergely Maróti, Carolina Falcón García, Oliver Lieleg, Ákos T. Kovács

**Affiliations:** Bacterial Interactions and Evolution Group, Department of Biotechnology and Biomedicine, Technical University of Denmark, Kgs Lyngby, Denmark; Terrestrial Biofilms Group, Institute of Microbiology, Friedrich Schiller University Jena, Jena, Germany; Seqomics Biotechnology Ltd, Mórahalom, Hungary; Institute of Plant Biology, Biological Research Centre, Hungarian Academy of Sciences, Szeged, Hungary; Department of Mechanical Engineering and Munich School of Bioengineering, Technical University of Munich, Garching, Germany

**Keywords:** experimental evolution, morphotype, pellicle, social interactions, biofilm matrix, *Bacillus subtilis*

## Abstract

Microbial biofilms are tightly packed, heterogeneous structures that serve as arenas for social interactions. Studies on Gram negative models reveal that during evolution in structured environments like biofilms, isogenic populations commonly diversify into phenotypically and genetically distinct variants. These variants can settle in alternative biofilm niches and develop new types of interactions that greatly influence population productivity. Here, we explore the evolutionary diversification of pellicle biofilms of the Gram positive, spore-forming bacterium *Bacillus subtilis*. We discover that - similarly to other species - *B. subtilis* diversifies into distinct colony variants. These variants dramatically differ in biofilm formation abilities and expression of biofilm-related genes. In addition, using a quantitative approach, we reveal striking differences in surface complexity and hydrophobicity of the evolved colony types. Interestingly, one of the morphotypes completely lost the ability of independent biofilm formation and evolved to hitchhike with other morphotypes with improved biofilm forming abilities. Genome comparison suggests that major phenotypic transformations between the morphotypes can be triggered by subtle genetic differences. Our work demonstrates how positive complementarity effects and exploitative interactions intertwine during evolutionary diversification in biofilms.

## INTRODUCTION

Rapid evolution of microbes constitutes a tremendous challenge to modern medicine, and at the same time, a privilege for microbial ecology. Due to large population sizes coupled with short generation times, bacterial adaptation can be observed in a course of days or months, allowing for its experimental investigation (Kawecki *et al.* 2012; Rosenzweig and Sherlock 2014; Martin *et al.* 2016). Experimental evolution studies continuously deepen our understanding of microbial adaptation, revealing common evolutionary scenarios such as genome reduction (Nilsson *et al.* 2005) or genome rearrangements (Martin *et al.* 2017), hypermutability (Flynn *et al.* 2016; Tenaillon *et al.* 2016), or diversification (Rainey and Travisano 1998; Poltak and Cooper 2011; Traverse *et al.* 2013; Koch *et al.* 2014; Flynn *et al.* 2016; Kim, Levy and Foster 2016). The last one, where microbes diversify into distinct variants (typically referred to as morphotypes as they are identified based on distinct colony morphology), appears to be very common, especially in structured environments which offer alternative niches varying in nutrient and oxygen content (Martin *et al.* 2016; Steenackers *et al.* 2016). Biofilms, where microbes grow in tightly packed assemblies, represent an example of such an environment.

Evolutionary diversification tends to improve biofilm productivity as newly emerged variants specialize in occupying different niches thereby reducing competition (Rainey and Travisano 1998; Poltak and Cooper 2011; Ellis *et al.* 2015; Flynn *et al.* 2016). An excellent example derives from a comprehensive study of an opportunistic pathogen, *Burkholderia cenocepacia,* which was allowed to evolve in a form of submerged biofilm subsequently assembling on a polystyrene bead floating in the liquid medium (Poltak and Cooper 2011). *B. cenocepacia* diversified into three distinct morphotypes, the two of which preferentially attached to the polystyrene surface, whereas the third mostly resided on the top of the biofilm (Ellis *et al.* 2015). The productivity of this ecotype mix was elevated due to the so called niche complementarity effect (Poltak and Cooper 2011; Ellis *et al.* 2015). Similar niche specialization was earlier observed in *Pseudomonas fluorescens* evolved in a static microcosm, where the population diversified into a ‘Wrinkly’ type colonizing the liquid-air interface, and ‘Smooth’ and ‘Fuzzy’ types residing at the bottom of the vessel or floating in the medium, respectively (Rainey and Travisano 1998).

Although evolutionary diversification is often associated with increased community productivity, interactions between the evolved variants can be complex and not necessarily positive. For example, *Pseudomonas aeruginosa* biofilms diversify into multiple variants differing in levels of key biofilm-stimulating messenger cyclic diguanosine monophosphate (c-di-GMP) (Flynn *et al.* 2016). These differences result in sequential surface colonization by the morphotypes, some of which remain in strong competition. In addition, subtle changes in c-di-GMP levels in one morphotype can dramatically shift the population structure, reducing the frequency of the c-di-GMP overproducer for the benefit of other variants (i.e. for the poor biofilm former) that, in turn, may decrease population productivity (Flynn *et al.* 2016). Analogous interplay takes place in *P. fluorescens* where production of cellulose by a surface colonizer, the ‘Wrinkly’, creates an opportunity for exploitative invasion by the ‘Smooth’ which alone cannot colonize the surface (Hammerschmidt *et al.* 2014). Invasion and, as a consequence, premature pellicle collapse happens because the ‘Wrinkly’ variant secretes a costly extracellular product, an acetylated cellulose (Spiers *et al.* 2003), that becomes vulnerable to exploitation by the non-producer. The two morphotypes coexist since each has an advantage when present at low frequency (Rainey and Travisano 1998).

Altogether, experimental studies on adaptation in biofilms seem to share certain scenarios: variants with distinct colony morphology readily emerge; the derived morphotypes differ in biofilm formation capabilities (Rainey and Travisano 1998; Poltak and Cooper 2011; Traverse *et al.* 2013; Leiman *et al.* 2014; Flynn *et al.* 2016); these differences result in a new type of spatial arrangement which reduces niche overlap and improves population productivity (Rainey and Travisano 1998; Poltak and Cooper 2011; Ellis *et al.* 2015; Flynn *et al.* 2016; Kim, Levy and Foster 2016). While the evolution in biofilms of medically-relevant Gram-negative microbes (*B. cenocepacia* and *Pseudomonas spp)* has been extensively studied, adaptive diversification in biofilms of Gram-positive bacteria has not received much attention. *Bacillus subtilis* is a wide spread, plant growth promoting spore-former, known from its ability to form complex pellicle biofilms at the liquid-air interface (Branda *et al.* 2001). Previous studies have already indicated that this bacterium readily diversifies into distinct colony types when cultivated under static or planktonic conditions (Leiman *et al.* 2014). Here, we explore the dynamics and ecological interactions during adaptive diversification in *B. subtilis* pellicle biofilms evolved in six parallel ecosystems.

When cultivated under static conditions, *B. subtilis* initially grows suspended, but increasing cell density results in decreasing oxygen concentration in bottom layers of the medium. Using aerotaxis, cells actively swim towards the liquid-air interface and colonize it in a form of densely packed pellicle (Hölscher *et al.* 2015). Pellicle formation requires secretion of extracellular matrix (ECM) comprising mainly exopolysaccharide EPS, fiber protein TasA and a hydrophobin called BslA. Lack of EPS or TasA prevents pellicle formation, but these matrix components can be shared by producers with the non-producers (Romero *et al.* 2011; van Gestel, Vlamakis and Kolter 2015; Martin *et al.* 2017). In contrast to rather slimy pellicles of *P. fluorescence* or *P. aeruginosa*, biofilms of *B. subtilis* yield a rigid and highly hydrophobic mat (Vlamakis *et al.* 2013; Arnaouteli, MacPhee and Stanley-Wall 2016). The non-wetting properties of those biofilms and pellicles are due to the extracellular matrix, with a major role of BslA (Kobayashi and Iwano 2012). The hydrophobicity of *B. subtilis* biofilms can be associated with the level of surface complexity (Werb *et al.* 2017). Comparable properties are observed for both colony and pellicle biofilms, and the ability to form wrinkly pellicles correlates with the ability to develop complex colonies (Romero *et al.* 2011; Shemesh and Chai 2013).

In this study we ask whether, likewise Gram negative species, *B. subtilis* can undergo reproducible evolutionary diversification, whether this diversification will involve easily quantifiable biofilm-related traits and finally, whether it will result in new types of social interactions.

## MATERIALS AND METHODS

### Strains and cultivation conditions

Supplementary Table S1 describes the strains used in this study and construction of their mutant derivatives. Strains were maintained in Lysogeny broth (LB) medium (LB-Lennox, Carl Roth; 10 g/L tryptone, 5 g/L yeast extract, and 5 g/L NaCl), while MSgg medium was used for biofilm induction (Branda *et al.* 2001).

### Experimental evolution and productivity assays

Experimental evolution was performed using the natural competent derivative of the undomesticated *B. subtilis* NCBI 3610, DK1042 strain (Konkol, Blair and Kearns 2013) grown in MSgg medium statically in a 24-well plate at 30°C for 48 hours in 6 parallel replicates. Mature pellicles were gently harvested from the surface of the liquid medium using plastic inoculating loop. First, the edge of the pellicle was gently pierced to partially detach the biofilm from the wall of the plastic well. Second, the pellicle was collected by rapidly moving the loop clockwise, constantly keeping the loop at the liquid-air interface and touching the wall/edge of the pellicle. This allowed harvesting of the entire biofilm, leaving the conditioned medium clear (optical density at 600nm~0). The material was then transferred to 2ml Eppendorf tube containing 1ml of 0.9% NaCl and 100μl of glass sand and vortexed vigorously for 60 seconds. This allowed efficient disruption of the material into single cells and small clumps, without the need for sonication which would highly increase the risk of contamination. Finally, the disrupted culture was reinoculated after 100× dilution. After the 5^th^,10^th^, 14^th^,19^th^, 24^th^, 29^th^ and 35^th^ pellicle transfers CFU/ml in the pellicles described here as pellicle productivity, were monitored and frozen (-80°C) stocks were preserved.

Single isolates representing the four different morphotypes were isolated from population 1. Productivity assays in mixed cultures containing different morphotypes were performed by mixing certain ratios of 100-fold diluted LB-pregrown cultures that were then incubated in static pellicle forming conditions for 48 hours. Prior each CFU assay, pellicles were sonicated according to a protocol optimized in our laboratory (Martin *et al.* 2017). Expected productivity was calculated by multiplying the frequency of a given morphotype in the initial inoculum by its productivity in monoculture (Poltak and Cooper 2011).

### Colony morphology assay

Colony morphologies were examined on LB and MSgg medium with 1.5% agar. The plates were dried under laminar airflow conditions for 20 min after solidifying. 2 μL of the overnight grown cultures were spotted on the plate, and the lids were closed once the spotted culture had dried. The plates were incubated at 30°C for 48 hours.

### Profilometric analysis

The surface profiles of colonies were obtained using a NanoFocus μsurf light profilometer (NanoFocus AG, Oberhausen, Germany). Randomly selected spots within colony center and colony periphery were scanned using 20× magnification resulting in images covering an 800×722 µm area. Missing data points were interpolated, and the scanned area was then evaluated with the software µsoft (Version 6.0, NanoFocus AG, Oberhausen, Germany) to obtain the developed interfacial area ratio (*Sdr* value). This metrological parameter determines the level of surface complexity (Werb *et al.* 2017): whereas *Sdr*=0 in the case of perfectly flat surfaces, the *Sdr* values are high for surfaces containing multiple grooves and valleys.

### Wetting

To test the wetting behaviour of colonies, a 10 μL water droplet was placed onto the colony center (and the colony periphery if the colony was large enough), and an image of the droplet were captured using a horizontal camera setup. Contact angles were extracted from this images using the “drop snake” plug-in in the software ImageJ.

### Spent media complementation assay

The supernatants (SN) were obtained from the ancestor strain and the Smooth strain grown under static conditions at 30°C. The SNs were sterilized using Milipore filters (0.2µm pore size) and mixed in 1:1 ratio with 2× MSgg resulting in MSgg+ancestor SN and MSgg+Smooth SN, respectively. Next the pellicle formation of the Smooth in presence of MSgg+ancestor SN was compared with the negative control, MSgg+Smooth SN.

### Microscopy/confocal laser scanning microscopy (CLSM)

Bright field images of whole pellicles and colonies were obtained with an Axio Zoom V16 stereomicroscope (Carl Zeiss, Germany) equipped with a Zeiss CL 9000 LED light source and an AxioCam MRm monochrome camera (Carl Zeiss, Germany). For single-cell level epifluorescence measurements, pellicles were harvested into 1.5 ml Eppendorf tubes containing 500 μl sterile MSgg base medium, vortexed for 90 seconds, mildly sonicated at 10% amplitude 12 seconds 1 second pause (Soniprep 150). 5 μl of sonicated cells was spotted on microscope slide covered with 0.8% agarose, covered with clean cover slip and examined under the fluorescence microscope (Olympus Bx51; 100× FASE Oil Objective; 98 ms exposure time).

Images were captured using the bright light and fluorescence light using the GFP filter. This procedure was done on two separate days with two biological replicates per day.

The pellicles were also analyzed using a confocal laser scanning microscope (LSM 780 equipped with an argon laser, Carl Zeiss, Germany) and Plan-Apochromat/1.4 Oil DIC M27 63× objective. Fluorescent reporter excitation was performed at 488 nm for green fluorescence and at 564 nm for red fluorescence, while the emitted fluorescence was recorded at 484–536 nm and 567–654 nm for GFP and mKate, respectively. To generate pellicle images, Z-stack series with 1 µm steps were acquired. Zen 2012 Software (Carl Zeiss, Germany) was used for both stereomicroscopy and CLSM image visualization.

### Genome resequencing and genome analysis

Genomic DNA of selected isolated strains was extracted using the EURx Bacterial and Yeast Genomic DNA Kit from cultures grown for 16 h. Paired-end fragment reads (2 × 150 nucleotides) were generated using an Illumina NextSeq sequencer. Primary data analysis (base-calling) was carried out with “bcl2fastq” software (v2.17.1.14, Illumina). All further analysis steps were done in CLC Genomics Workbench Tool 9.5.1. Reads were quality-trimmed using an error probability of 0.05 (Q13) as the threshold. In addition, the first ten bases of each read were removed. Reads that displayed ≥80% similarity to the reference over ≥80% of their read lengths were used in mapping. Non-specific reads were randomly placed to one of their possible genomic locations. Quality-based SNP and small In/Del variant calling was carried out requiring ≥8× read coverage with ≥25% variant frequency. Only variants supported by good quality bases (Q ≥ 20) were taken into account and only when they were supported by evidence from both DNA strands. Data on mutation frequencies for each strain sequenced in the study are provided in Supplementary Table S2.

## RESULTS

### Evolution of pellicle biofilms of *B. subtilis* involves rapid diversification into four distinct morphotypes

To examine whether evolutionary diversification can be observed in *B. subtilis* NCBI 3610 pellicle biofilms, we allowed 6 parallel pellicle populations to evolve for over ca. 200 generations (see methods). Pellicles were visually accessed after each subsequent transfer, and their productivities (total cell number/ml) were assessed every 5^th^ transfer. All pellicles appeared robust throughout the experiment preserving a thick and wrinkly structure similar to the ancestral strain. The productivity assay suggested that, with certain exceptions (e.g. #35), CFU/ml of all biofilm populations increased gradually thorough the evolution experiment (Figure 1A). The significant increase in productivity was confirmed by regression analysis when considering all replicates regardless of the timeframe analyzed (0-24# or 0-35#) (Supplementry Table S3). When looking at individual ecosystems, the statistical significance could only be confirmed for replicate 4 (0-24#; p<0.02), while replicate 6 served as an outlier with dramatic productivity drop at 35#, resulting in its lower final as compared to initial productivity (0-35#; p<0.99). On the same line, correlation analysis revealed strong synchrony in productivity changes across all populations, except for the population 6, which exhibited the lowest values of Pearson’s correlation coefficient when compared to all remaining populations (Supplementary Table S4).

**Figure 1.**
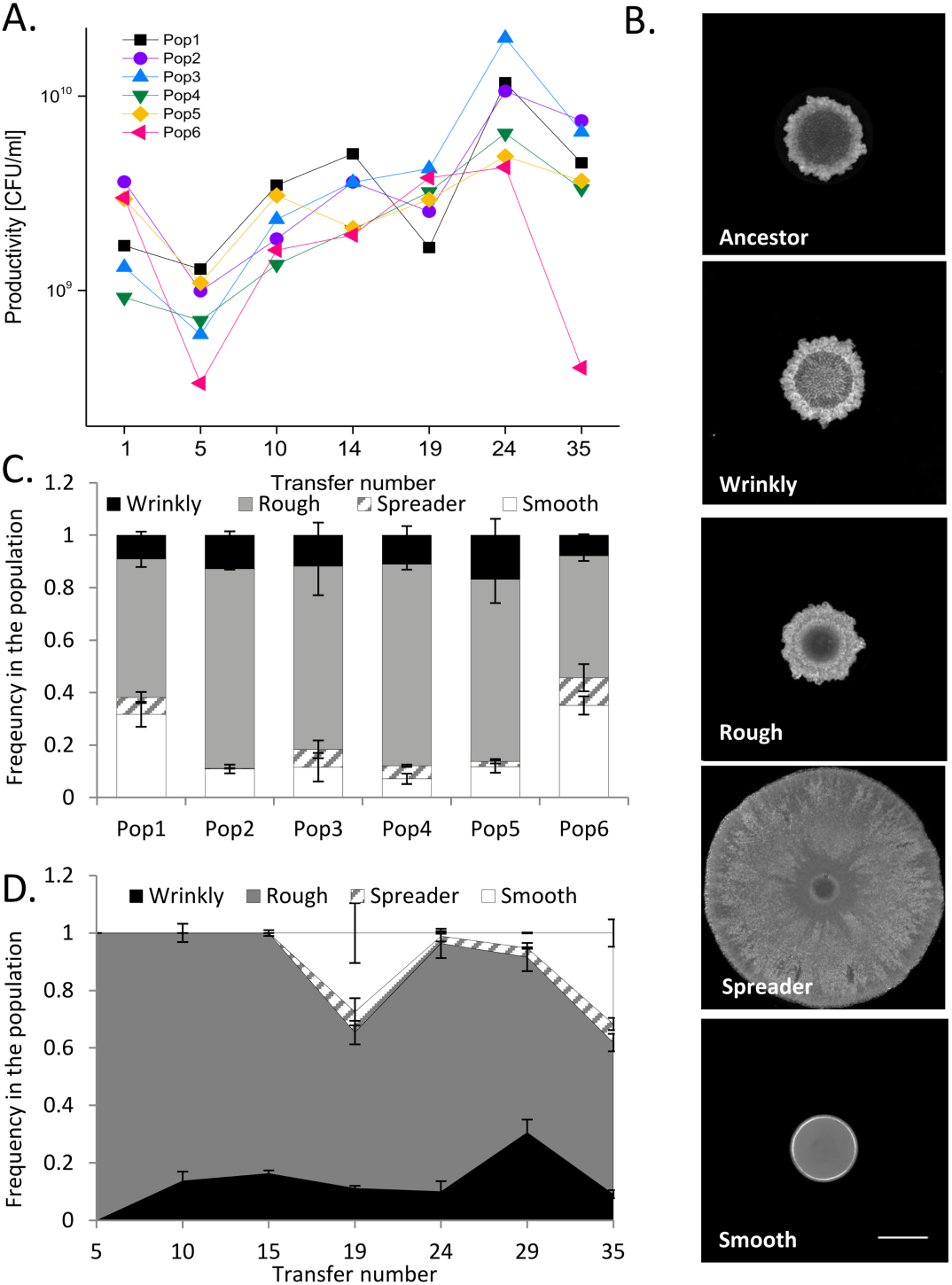
Productivity changes and diversification in *B. subtilis* pellicles. **(A)** Biofilm productivities were systematically monitored (as CFU/ml) during ongoing evolution experiment. Productivities were assessed every 5^th^ or 4^th^ transfer, with an exception of the two final time points, where sampling gap was larger (11 transfers). **(B)** Distinct colony morphologies of the evolved variants were especially pronounced on solid LB medium. The ancestor *B. subtilis* NCBI 3610 and four morphotypes isolated from population 1 and were spotted on LB agar (1.5%) and imaged after 48h of incubation at 30°C. The names (Wrinkly, Rough, Spreader and Smooth) were assigned arbitrarily, based on colony appearance. Scale bar represents 5mm. **(C)** Final (35^th^ transfer) relative frequencies of different morphotypes (Wrinkly, Rough, Spreader and Smooth) in all 6 parallel evolved populations (for each population n=3). Data represent the mean and error bars represent standard error. **(D)** Relative frequencies of different morphotypes in subsequent evolutionary time points in population 1 (n=3). For panel C and D: data represent the mean and error bars represent standard error.

In all 6 populations, four distinct colony types (morphotypes) could easily be detected in CFU assay. A representative example from each morphotype was isolated from population 1 and stored as pure culture stock for further studies. To better assess differences between the morphotypes, a colony spotting assay was performed (see methods) on two alternative media types: rich LB medium (Fig. 1B) and minimal biofilm-promoting MSgg medium (Branda *et al.* 2001) (Supplementary Fig. S1A). Inspired by the striking difference in appearance of the morphotypes on LB medium, we introduced the following names: Wrinkly - displaying an increased complexity in the colony center; Rough - similar to the ancestor; Spreader - showing dramatically increased colony expansion; and Smooth – exhibiting a very flat surface similar to certain biofilm mutants (Romero *et al.* 2010) (Fig. 1B).

To determine the final frequencies of these morphotypes in 6 parallel evolved populations, small fractions of the corresponding −80°C stocks were serially diluted and plated on LB-agar. The frequency of the Wrinkly morphotype was similar in all populations ranging from 8% in population 6 up to 17% in population 5 (Fig. 1C) (Supplementary Table S5). The Rough derivative was the most abundant morphotype in all populations with the highest relative abundance in populations 2 and 4 (76% and 77%, respectively) and least dominant in population 6 (47%). The Spreader showed a maximum of 10% in population 6, while it was barely detected in population 2 (0.2%). Finally, the Smooth variant showed the highest frequency in populations 6 and 1 accounting for 35% and 31% of the pellicles, respectively (Fig. 1C). Statistical analysis revealed that in these two populations the frequency of Smooth was significantly higher than in populations 2-5 (Supplementary Table S5). High frequency of Smooth in populations 1 and 6 was coupled with lower frequency of Rough morphotype in these populations (Fig. 1C, Supplementary Table S5).

We were next interested in the evolutionary history of diversification in pellicles. To assess the subsequent emergence of the detected four morphotypes during the experimental evolution, frozen pellicle stocks of population 1 from 7 chronological time-points (i.e., from every 5^th^ transfer) were re-examined. All colonies obtained from the 5^th^ transfer resembled the Rough morphotype; however, it is important to note that the ancestor and the Rough variant are very similar (Fig. 1D). The Wrinkly variant was first observed in the 10^th^ transfer and its frequency remained relatively constant until the 35^th^ transfer. The Spreader and Smooth variants both emerged around the 19^th^ pellicle. Whereas the frequency of the Spreader variant remained very stable throughout the evolution, the frequency of the Smooth morphotype was oscillating: it dropped to nearly zero in the 24^th^ pellicle but it recovered to over 30% at later evolutionary time points (Fig. 1D). Interestingly, the frequency of the Smooth variant inversely matched with the productivity pattern throughout the evolutionary history of population 1 (Fig. 1A): transfers where the productivity dropped displayed an increase in frequency of the Smooth morphotype (Fig. 1D) (linear regression analysis gave negative slope with r^2^ = 0.32; p<0.06). Similarly, population 6 had the lowest final productivity but the highest frequency of the Smooth morphotype (Fig. 1C) (negative slope with r^2^= 0.26; p<0.03).

### Evolved morphotypes develop colonies with distinct surface complexity and hydrophobicity

Next, we attempted to quantitatively describe differences between the observed morphotypes and examined the microscopic surface profiles of the pellicle colonies. Surface profilometry was performed for all morphotypes and the ancestor, both on LB and MSgg medium (Fig. 2, Supplementary Fig. S1). Although the results were affected by the media type, certain pronounced differences between the morphotypes were media-independent. The Smooth variant showed a lack of surface complexity on both media types which was reflected by a *Sdr* value of practically 0 (see methods). On MSgg medium, the Ancestor and Rough morphotypes showed the highest surface complexity which was about 4-fold higher as compared to the colonies of the Wrinkly and Spreader variants (Fig. 2A-B). On LB medium, all morphotypes depicted lower surface complexity as compared to the ancestor, with higher *Sdr* values for the Wrinkly and Rough derivatives (Supplementary Fig. S1B).

**Figure 2.**
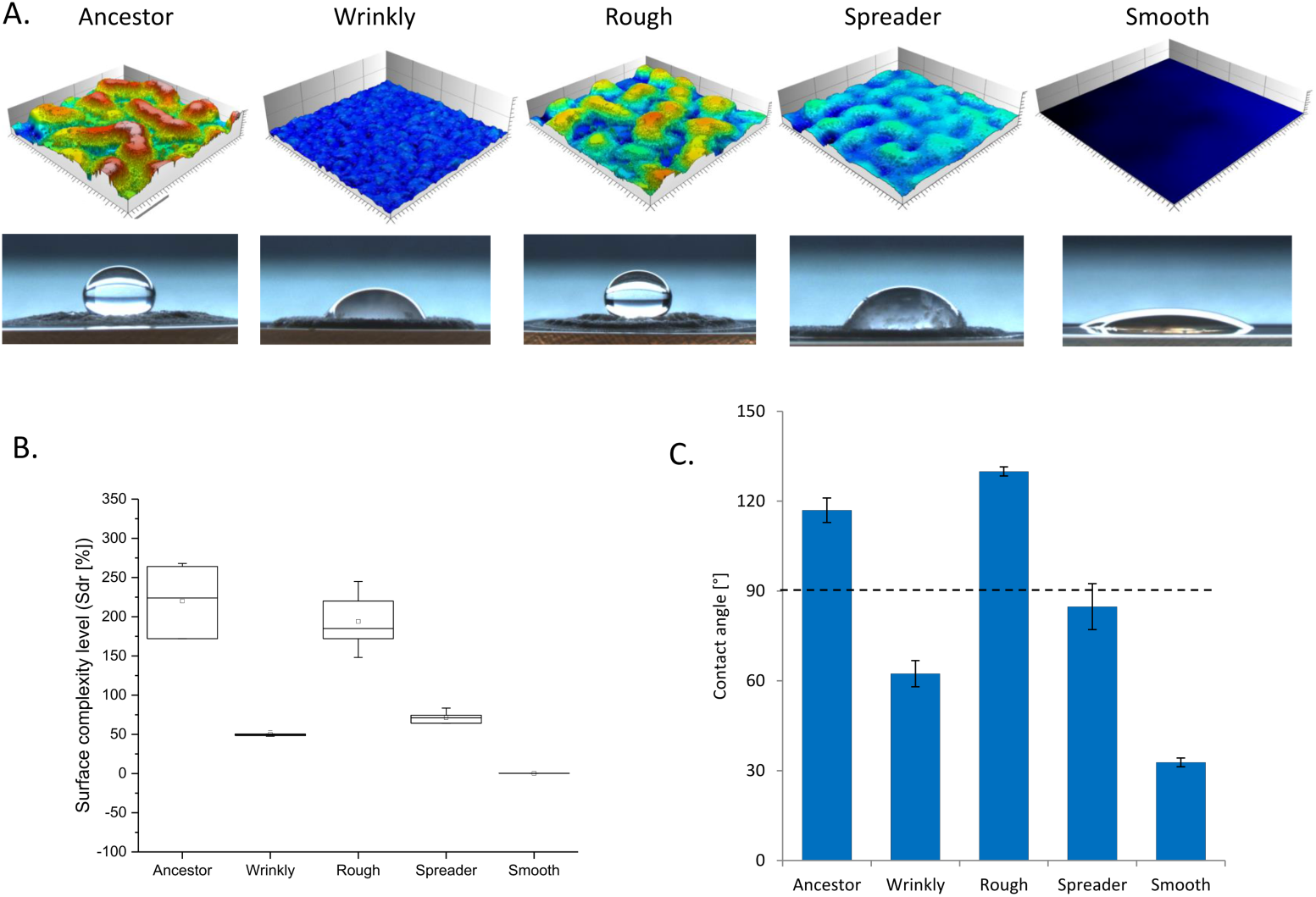
Quantitative characterization of colony features displayed by the evolved morphotypes on biofilm-promoting MSgg medium. **(A)** Surface topologies of colony center of the ancestor and evolved morphotypes were acquired using light profilometry. 3D images of the surface topology were generated using the software μsurf (see methods) showing topological features with a standard color scale-where dark blue represents the lowest and white represents the highest features. Colonies were grown for 48h at 30°C. Scale bar represents 200µm. Below: image of 10µl water droplet spotted at the corresponding colony center. **(B)** Developed interfacial area *Sdr* calculated for colony center based on surface topology (some samples n=6, while others n=5). Boxes represent Q1–Q3, lines represent the median, and bars span from max to min. **(C)** Contact angles calculated for water spotted in colony center (n=3). Dashed line represents contractual hydrophobicity cut-off, separating the hydrophilic (below the line) from hydrophobic (above) surfaces. Data points represent mean and error bars represent standard error.

Analyses of surface profiles of expanding colony edges showed that, when cultivated on MSgg medium, only the Spreader variant exhibited lower surface complexity at the edge than the ancestor; in contrast, on LB, all strains showed lower *Sdr* values on the edge as compared to the ancestor (Supplementary Fig. S2).

As previous studies suggested that the complexity of the biofilm surface might directly correlate with its hydrophobicity (Werb *et al.* 2017), we additionally performed wetting studies using the four morphotypes (see methods). In line with those previous results, the Smooth variant was lacking typical biofilm hydrophobicity and acted completely hydrophilic on both LB and MSgg media (Fig. 2A,C; Supplementary Fig. S1). When grown on Msgg medium, the Rough morphotype showed a non-wetting behavior similar to the ancestor, but not on LB; here, the Wrinkly and Rough variants both showed hydrophilic properties (Fig. 2A,C; Supplementary Fig. S1). Interestingly, while the Spreader morphotype was generally hydrophilic, its expanding edge showed highly hydrophobic properties outperforming those of the ancestor strain (Supplementary Fig. S1C).

Overall, these studies revealed that evolutionary diversification led to clear and measurable differences in the colony surface properties with correspondingly different wetting properties.

### Morphotypes display distinct levels of matrix gene expression

Since previous experiments revealed dramatic differences in colony surface complexity and wetting behavior, we further hypothesized that these differences could be linked to different levels of extracellular matrix (ECM) production by the morphotypes (Kobayashi and Iwano 2012; Kesel *et al.* 2016). In order to test this, the matrix-genes reporters P_*tapA*_-*gfp* and P_*eps*_-*gfp* were introduced into the four morphotypes and the ancestor strain, and changes in matrix genes expression were determined (Fig. 3A, Supplementary Fig. S3, Supplementary Fig. S4).

**Figure 3.**
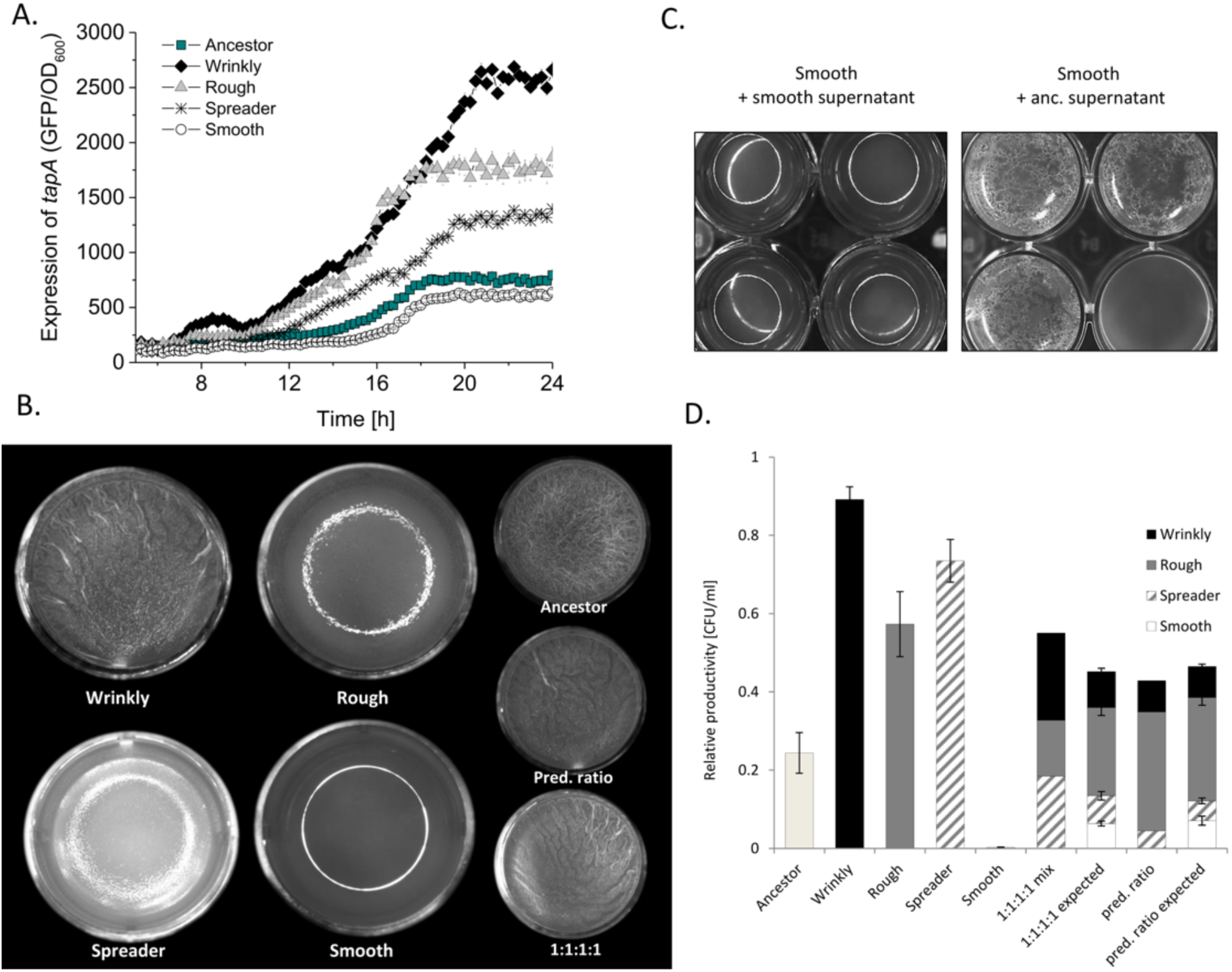
Biofilm genes expression and pellicle productivities in monocultures and in the mix. **(A)** Expression of *tapA* gene in the ancestor and all four morphotypes was compared using corresponding strains carrying P_*tapA*_-gfp reporter fusion. The experiment was terminated after 24 hours due to clumping of cells which interferes with reliable OD and fluorescence signal reads (n=8). Data points represent mean and error bars represent standard error. **(B)** Pellicle morphology developed by the ancestor, all morphotypes grown in monocultures and two types of morphotype mixes: equal frequencies of morphotypes in initial inoculum (1:1:1:1); predicted frequencies of morphotypes in initial inoculum (0.9: 5.3:0.6:3.2) that are matching their original ratios in evolved pellicle (see Fig. 1B,C). Pellicle were grown for 48h at 30°C. **(C)** Effect of the spent media produced by the ancestor on pellicle formation by the Smooth morphotype. Four wells were imaged to represent the biological variation of the assay. Well diameter is 1.5 cm. **(D)** Productivities of the ancestor, Wrinkly, Rough, Spreader and Smooth morphotypes, grown in monoculture and in mixes. Expected productivity was calculated as the product of the proportion of each morphotype in the initial population and its yield (CFU per ml) in monoculture (Loreau and Hector, 2001; Poltak & Cooper). Observed productivity is the total yield of the mixed community in the experimental environment (for monocultures n=9; for mixes n=12). Data points represent mean and error bars represent standard error.

Fluorescence images of colonies developed by P_*tapA*_-*gfp* and P_*eps*_-*gfp* labelled strains suggested the lowest expression of both matrix genes in the Smooth variant, moderate expression (comparable to the ancestor) in the Spreader variant and increased expression in case of the Rough and Wrinkly variants (Supplementary Fig. S3, Supplementary Fig. S4). To confirm this observation, the P_*tapA*_-*gfp* fluorescence was quantified during planktonic growth in all morphotypes and the ancestor (Fig. 3A). As expected, the Wrinkly and Rough morphotypes showed dramatic increase in *tapA* expression as compared to the ancestor. The detected fluorescence level from the P_*tapA*_-*gfp* construct in the Spreader variant was also increased. Finally, the Smooth variant showed decreased P_*tapA*_-*gfp* fluorescence as compared to the ancestor indicating lower levels of matrix genes expression (Fig. 3A). We also compared the expression of *tapA* in the ancestor and evolved morphotypes at single-cell level (Supplementary Fig. S5). While Wrinkly, Rough and Spreader preserved phenotypic heterogeneity pattern of matrix genes expression similar to the ancestor, *tapA* expressing cells could barely be detected in the Smooth population (Supplementary Fig. S5).

As expected based on colony morphology and low levels of matrix genes expression, the Smooth morphotype was not able to form a pellicle biofilm when cultivated solitarily (Fig. 3B). Pellicles formed by the Wrinkly variant resembled the ancestral pellicle, but those formed by the Rough and Spreader variants showed a less wrinkled and shinier surface structure as compared to the ancestor (Fig. 3B). Pellicles formed by the mix of all 4 morphotypes showed a typical wrinkly structure with a matte surface similar to the ancestor, regardless whether the initial ratio was 1:1:1:1 or the derivatives were mixed according to ratios established in the last stage of evolution, i.e. pellicle 35 (Fig. 3B).

Since the Smooth morphotype could not develop a pellicle but was detectable in the mixed population (Fig. 1), we tested whether biofilm formation by the Smooth variant can be complemented by the supernatant produced by the pellicle-forming ancestor that secretes a functional ECM. Although a robust pellicle could not be completely restored, the Smooth morphotype displayed improved surface colonization when grown in the presence of the ancestors spent medium (Fig. 3C), but not when its own supernatant was applied. Therefore, the Smooth variant can benefit from extracellular substances released by other non-defective strains and thereby co-colonize the liquid-air interface.

### Non-matrix producing morphotype exploits matrix-producing variants

Finally, we inspected the impact of diversity on individual and group performance by the morphotypes by comparing the productivity of monoculture pellicles with the mixed pellicles in addition to determining the frequency of each morphotype in the mix (as previously performed by Poltak and Cooper 2011). All morphotypes except the Smooth variant performed significantly better than the ancestor. The Rough and the Spreader morphotypes displayed over 2-or 3-fold higher productivity, respectively, while the Wrinkly variant showed the highest productivity (Fig. 3D). The effect of diversity was assessed using two types of mixes with alternative inoculation strategies: (A) using equal initial frequencies of all morphotypes (1:1:1:1) or (B) inoculating with frequencies observed in the evolved pellicle (i.e., Wrinkly 9%, Rough 53%, Spreader 6%, and Smooth 32%). The expected productivities in the mix were calculated as a product of their initial frequencies in the mix and the monoculture yield (as previously defined in Poltak and Cooper 2011). First, we discovered that the Wrinkly, Rough and Spreader monoculture yields were higher than the yields of the mixes (Fig.3C). Interestingly, the net biodiversity effect on the yield was dependent on the inoculation strategy. Specifically, in mix A, the observed yield of the mix was lower than the predicted productivity indicating interference effects. The productivity changes of the Wrinkly, Rough and Spreader variants in the mix were moderate (0.41-fold, 1.57-fold and 0.39-fold, respectively), but the productivity of biofilm-defective Smooth showed a dramatic 98.2-fold increase (Fig. 3D). In mix B (containing altered frequencies), the individual productivity changes of the Wrinkly, Rough and Spreader variants were minimal (0.99-fold, 0.87-fold and 1.15-fold, respectively), while the amplified abundance of the Smooth variant was slightly reduced to 86.6-fold (Fig. 3D), overall resulting in a slightly positive effect on biodiversity and community yield. The calculated effects of selection and complementarity revealed negative selection effects in both mixes (A: −13.4, B: −14.82), balanced by positive complementary effects (A: 13.3, B: 14.86), and resulting in slightly positive net effects in mix B.

It is interesting to note that the final frequencies of the morphotypes in mix A and B were nearly identical despite the differences in the initial inocula. We hypothesized that the robust structure of the community can be governed by certain assortment patterns of the morphotypes in the pellicle. To test whether morphotypes are non-randomly distributed in the pellicles, we fluorescently labelled all morphotypes and evaluated the assortment of each morphotype in a pairwise assay (Fig. 4, Supplementary Fig. S5). Confocal laser scanning microscopy analysis revealed that the Smooth variant was mostly localized at the bottom layer of the pellicle probably attaching to the matrix-producing strains (Fig. 4, Supplementary Fig. S5). Both the Spreader and the Wrinkly variants were present across the whole intersection of the pellicle, although at lower frequencies as compared to the Rough variant (which is in line with cell count results, see Fig. 3D) (Fig. 4, Supplementary Fig. S5). Although technical limitations did not allow us to visualize all morphotypes in the pellicle simultaneously, the pairwise analysis suggests that the Rough, Wrinkly and Spreader variants share the same niche, while the Smooth variant is mostly localized in the lower, medium proximal sections of the pellicle (Fig. 4, Supplementary Fig. S5).

**Figure 4.**
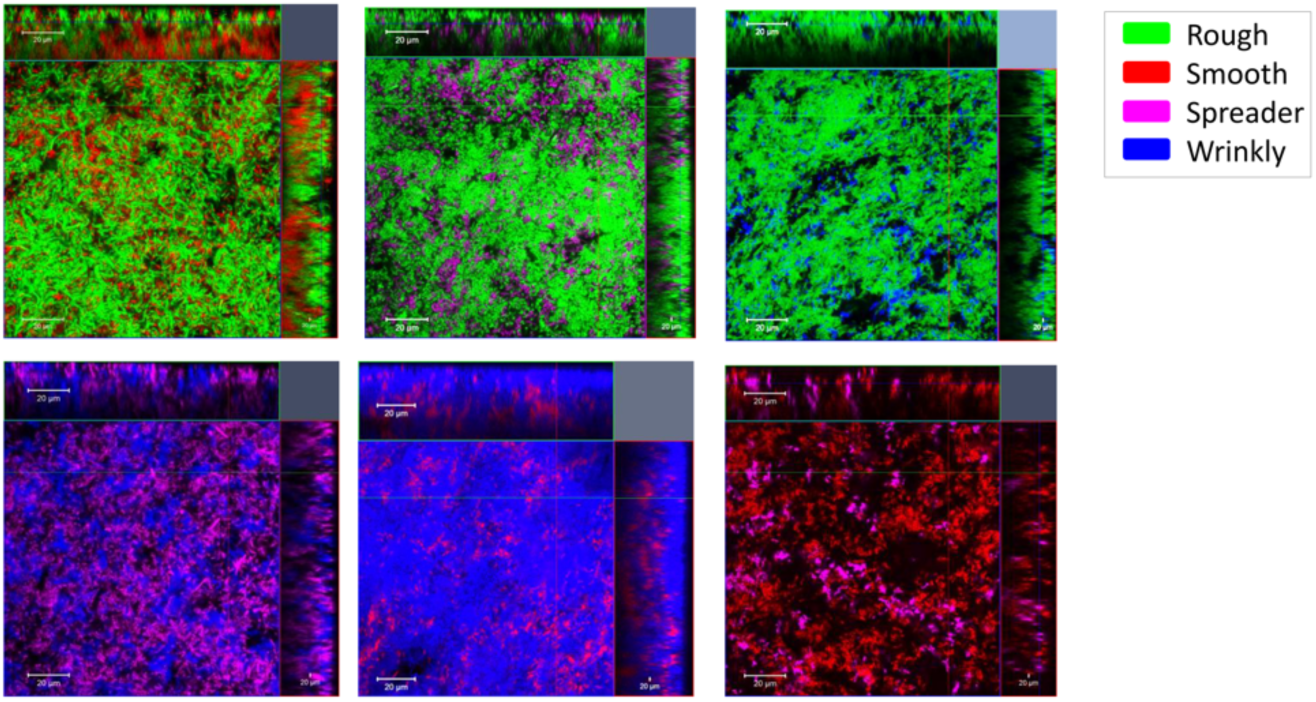
Spatial assortment of morphotypes in pellicle. Confocal microscopy images of pellicle biofilms formed by the mix of four morphotypes were taken. Each time only 2 selected morphotypes were labelled with constitutive fluorescent reporters and could be visualized in the pellicle. Morphotypes were mixed in ratios 0.9:5.3:0.6:3.2 ratios and pellicle were allowed to form for 48h at 30°C. To simplify image comparison, each morphotype was artificially labelled with the same assigned color (regardless on actual fluorescent marker used) across all images: Rough – green, Smooth – red, Spreader - purple, Wrinkly - blue. The actual mixed cultures were as follows: Rough_mKate_ + Smooth_GFP_ (+Wrinkly and Spreader unlabeled), Rough_GFP_ + Spreader_mKate_ (+Wrinkly and Smooth unlabeled), Rough_GFP_ + Wrinkly_mKate_ (+Spreader and Smooth unlabeled), Spreader_GFP_ + Wrinkly_mKate_ (+Rough and Smooth unlabeled), Smooth_GFP_ + Wrinkly_mKate_ (+Rough and Spreader unlabeled), Smooth_GFP_ + Spreader_mKate_ (+Rough and Wrinkly unlabeled).

### Diversification involves mutations related to biofilm regulation, motility and anaerobic respiration pathways

To understand the genetic bases of diversification and interactions between the morphotypes, the genomes of the Wrinkly, Rough, Spreader and Smooth variants isolated from population 1 were sequenced. In addition, 6 randomly picked isolates from other parallel populations were selected for sequencing (Table 1). The genome of the ancestor was also re-sequenced to screen for single SNPs that could emerge before the evolution experiment during standard stock preparation.

Colony morphology analysis later revealed that the additional strains 2A, 2B and 3C represented the Wrinkly morphotypes, while 1B, 4B and 6C represented the Rough morphotypes (Supplementary Fig. 6A). All isolates contained over 400 mutations of which 386 were overlapping and were therefore not subjected to further analysis, since they probably were linked to general adaptation to laboratory conditions or specifically to the medium. 58 non-overlapping mutations were detected in total, most of which resulted in amino-acid substitution or a frame shift (22 and 15, respectively). Remaining mutations were found in non-coding regions (11), rRNA (5) or resulted in synonymous substitutions (5). Certain mutations detected in non-coding regions (NC) and rRNA reproduced across representatives of different morphotypes, suggesting their early occurrence in evolution (Table1). On the other hand certain NC mutations were unique for the Smooth (2) or Spreader (1), and could indicate unknown regulatory role of those regions (Table 1). The most common mutation (detected in 7 out of 8 sequenced isolates) was synonymous substitution in *ppsD* gene which could be part of codon optimization process. Remaining mutations detected in coding regions could be divided into 6 categories: mutations detected in Wrinkly, Rough and Spreader strains, mutations detected in both, Wrinkly and Rough strains, mutations present solely in the Wrinkly strain, mutations exclusive to the Rough strain, mutations shared between the Smooth and the Spreader strain, and finally a single mutation present only in the Spreader strain (Table 1). There were 2 mutations that placed in the first category: Val66Leu substitution in the product of *qoxA* gene that encoding for quinol oxidase, and a synonymous substitution in *pyrP* (Table 1).

All Rough and all Wrinkly strains contained a substitution in HemAT, a soluble chemotactic receptor playing a crucial role in pellicle formation (Hölscher *et al.* 2015). The substitution was either Phe137Leu in case of Wrinkly 3C or Leu140Pro in other Wrinkly and Rough representatives (Table 1). Certain genes like *srfAA, pyrP* or *iolF,* encoding for surfactin synthetase, uracil permease and D-chiro-inositol transport protein, respectively, were reproducibly mutated in different Wrinkly/Rough strains but at different positions (Table 1).

All Wrinkly isolates shared a mutation in the *kinA* gene which is directly linked to the matrix master regulator Spo0A (Stephenson and Hoch 2002). In addition, all but one Wrinkly representative had non-synonymous substitution in *yogA* (unknown function) and frame shift mutation in *coaX* involved in Coenzyme A biosynthesis (Table 1). Mutations unique for the Rough group were rather scattered through different strains, but non-synonymous substitution in *polC* encoding for DNA polymerase III were found in two different strains at different positions of this gene (Table 1).

Both the Spreader and the Smooth strain exhibited a substitution in the *rex* gene which may play a role in their adaptation to the limited oxygen availability in the bottom layers of the pellicle. Interestingly, they also shared a substitution in the flagella-related gene *fliY*, but the Spreader strain additionally carries a frame shift mutation in the GMP-synthesis gene *guaA*. We examined the effects of *rex* and *guaA* deletion on pellicle and colony morphology, but neither resulted in a phenotype that would resemble that of Spreader or Smooth pellicles (Supplementary Fig. S6B) indicating that at least two different SNPs are needed to recreate the evolved morphotypes (assuming no effects of SNPs present in non-coding regions). The mutation pattern (Table 1) and the simultaneous emergence of Spreader and Smooth morphotypes during the evolution suggest that the Smooth variant could be the ancestral form of the Spreader variant. The Wrinkly morphotype likely evolved from the Rough variant, since it emerged after the Rough morphotype (Fig. 1C) and it contained mutations that overlap with those found in the Rough variant in addition to subsequent SNPs.

## DISCUSSION

Several studies have already proved that diversification is a rapid, general and significant process in microbial evolution. Our work fills the gap of equivalent knowledge on a non-pathogenic and biotechnologically relevant Gram-positive model. We reveal, that diversification pattern during *B. subtilis* pellicle evolution with respect to colony types that emerge from a common ancestor, reproduced in all parallel ecosystems. Although the detailed evolutionary history was examined only in 1 out of 6 parallel populations, the synchrony in productivity changes throughout the evolutionary time and similarities in the final frequencies of morphotypes across all 6 populations point towards strong parallelism of evolutionary events occurring in independent pellicles. We employed a novel quantitative approach to describe the evolved morphotypes and to confirm that they were different from each other and from their common ancestor. Surface profilometry combined with wetting studies were especially helpful in the case of early morphotypes (Rough or Wrinkly): whereas they still resembled the ancestral colony in terms of macro-scale morphology, remarkable differences in microstructure and hydrophobicity were revealed. Recent work indicated that subtle differences in colony microstructure could be of profound importance for colony wetting behavior (Arnaouteli *et al.* 2017; Werb *et al.* 2017) and, consequently, for its resistance to antimicrobials (Arnaouteli *et al.* 2017). However, recent findings clearly show that complex surface topology is not sufficient to maintain non-wetting behavior of *B. subtilis* colonies (Arnaouteli *et al.* 2017). To a certain extent, the measured surface complexity/hydrophobicity levels were positively correlated with the levels of matrix genes expression, but this correlation was not perfect. For instance, the Wrinkly variant showed a strongly increased expression of the *tapA* operon and yet a decreased surface complexity and hydrophobicity as compared to the ancestor. This is in line with a recent survey of domesticated *B. subtilis* 168 variants where the authors also found a certain mismatch between the colony surface complexity (at macroscale) and the level of biofilm-genes expression (Gallegos-Monterrosa, Mhatre and Kovács 2016). As a complex colony structure is affected by localized cell death (Asally *et al.* 2012), changes in cell viability in evolved morphotypes could potentially balance the effects of increased EPS production levels.

Similar to previous studies, *B. subtilis* evolved into variants with improved biofilm productivities (Rough, Wrinkly and Spreader); however, a quantitative analysis revealed that some of them lost the important surface properties of the ancestor (Wrinkly and Spreader), presumably pointing towards quantity vs quality trade-offs. Remarkably, one of the morphotypes (Smooth) completely lost the ability to form a pellicle, and it could only reside in such biofilms encompassing also the other variants. Moreover, the evolved non-producer (Smooth) was the only morphotype gaining a strong individual benefit from being part of the mixed population regardless of its initial frequency in the inoculum. The minor complementarity effects that were estimated when comparing the expected vs obtained mix productivity *in toto* may derive from the fact that the Smooth variant mostly resides in the bottom layers of the pellicle. This might reduce its competition with the biofilm-forming morphotypes due to restricted oxygen availability. The interaction between *B. subtilis* Smooth and other variants resembles the interplay between ‘Smooth’ and ‘Wrinkly’ morphotypes of *P. fluorescens* (Rainey and Travisano 1998), with the particular difference that, here, the “defector” evolves *de novo*. We believe that the success of the Smooth variant could depend on early adaptive events in the population, specifically the rise of Rough and Wrinkly variants that both showed a tremendous increase in EPS production. Still, the diversity within matrix-producers (in this case Spreader, Rough and Wrinkly) can actually limit the spread of matrix-non-producers (Brockhurst *et al.* 2006).

Such non-producers were not identified in evolving *B. cenocepacia* or *P. aeruginosa* biofilms. Possibly, submerged biofilms of *B. cenocepacia* or *P. aeruginosa* may strongly select for attachment and stratification of individual cells (Xavier and Foster 2007), while in floating biofilms of *B. subtilis* and *P. fluorescens*, cells can partially rely on medium diluted EPS. Although we did not unravel the molecular evolution pattern, certain mutations discovered in given morphotypes represent promising targets for future studies. For example, Rough and Wrinkly carried a Leu140Pro or Phe137Leu substitutions in HemAT, which is a key oxygen receptor important for pellicle formation (Hölscher *et al.* 2015). Since Phe137 is directly at heme-biding side and a substitution to Leu very likely alters the function of the receptor. Similar is the case for Leu140, part of the α-helix and only 3 amino acids away from the heme-binding side, where a non-conserved substitution to Proline likely influences the function of the receptor. Non-synonymous substitutions in *srfAA* detected in certain Wrinkly and Rough isolates could alter the function of surfactin synthetase (especially the non-conserved Ile2983Asn detected in Wrinkly 1, Rough 1 and 4B) what could influence the amounts of produced surfactin. As surfactin is believed affected matrix genes expression as a paracrine signaling molecule (López *et al.* 2009), it might have an impact the *eps/tapA* expression levels in those morphotypes. Both Smooth and Spreader strains share mutations in the *rex* and *fliY* gene, which could indicate their adaptation to the oxygen-poor bottom layers of biofilms and changes in motility, respectively. In addition, another substitution in *rex* was also found in Rough 6C and other genes linked to anaerobic respiration (*nirB*) or chemotaxis (*mcpC, cheA*) were also altered in selected Wrinkly and Rough isolates (Table 1).

Interestingly, all but two NC mutations present in the Smooth overlap with the Spreader, which contains 3 additional unique mutations only one of which (Thr151 fs in *guaA*) is positioned in a coding region. Presumably, supplementing Smooth genetic background with a *guaA* mutation might result in secondary effects that transform the Smooth into a Spreader phenotype. As c-di-GMP plays an important role in motility arrest (Chen *et al.* 2012a), mutation in GMP synthetize GuaA probably reduces the level of c-di-GMP possibly enhancing motility.

It is difficult to point towards the mutations that are responsible for Rough and Wrinkly morphotype. Rough 1B carries the lowest amount of mutations where only *qoxA* (related to respiration), *hemAT*, *murG* (peptidoglycan synthesis) and *pyrP* were identified to carry non-synonymous substitutions/frame shifts. Unless Rough resulted from changes in unknown regulatory elements, those 4 genes could be potential targets of molecular evolution. Morphogenesis of Wrinkly could require additional changes in *kinA*.

Variations in biofilm formation within laboratory *B. subtilis* strains are commonly associated with the domestication problem, that occurs during cultivation of a bacterium in rich LB medium (McLoon *et al.* 2011; Leiman *et al.* 2014; Gallegos-Monterrosa, Mhatre and Kovács 2016). Indeed, biofilm-altered variants of *B. subtilis* could be experimentally evolved in static and liquid LB, mostly carrying mutations in the biofilm regulator *sinR* (Leiman *et al.* 2014). Here, we showed that biofilm non-producers readily evolve also in minimal medium under biofilm selective conditions indicating they can also be part of a normal eco-evolutionary process, rather than just a product of domestication. We still know relatively little about the natural diversity of biofilm formation within *B. subtilis.* Such information would be of great importance taking into account biocontrol properties of the species which rely on biofilm formation capability (Chen *et al.* 2012b).

We here showed that evolution in pellicle biofilms may give rise to variants with variable biofilm phenotypes including biofilm non-producers that rely on ‘upgraded’ pellicle formers. Our work reveals that, in addition to positive niche opportunities, spatially structured environments provide a platform where positive interactions intertwine with exploitation.

